# *foxe1* mutant zebrafish show indications of a hypothyroid phenotype and increased sensitivity to ethanol for craniofacial malformations

**DOI:** 10.1101/2024.01.31.578204

**Authors:** Sophie T Raterman, Frank A D T G Wagener, Jan Zethof, Vincent Cuijpers, Peter H M Klaren, Juriaan R Metz, Johannes W. Von den Hoff

## Abstract

FOXE1 mutations in humans are associated with Bamforth-Lazarus syndrome, characterized by cleft palate and hypothyroidism. Moreover, polymorphisms of FOXE1 are implicated in non-syndromic cleft palate. Much uncertainty still exists about the function of transcription factor FOXE1 in development. To address this, we have previously developed a *foxe1* mutant zebrafish demonstrating mineralization defects in larvae. In the present study, we further investigate the thyroid status and skeletal phenotype of adult *foxe1* mutants. Compared to wild type controls, mutant fish have increased expression of hypothalamic *tshβ*, and hepatic *dio1* and *dio2*. In plasma we found higher circulating Mg levels; together these findings are indicative of hypothyroidism. We further observed mineralization defects in scales, likely due to enhanced osteoclast activity as measured by increased expression levels of the markers *tracp, ctsk* and *rankl*. Gene-environment interactions in the etiology of FOXE1-related craniofacial abnormalities remain elusive, which prompts the need for models to investigate genotype-phenotype associations. We here investigated whether ethanol exposure increases the risk of developing craniofacial malformations in *foxe1* mutant larvae that we compared to wild types. We found in ethanol-exposed mutants an increased incidence of developmental malformations and marked changes in gene expression patterns of cartilage markers (*sox9a*), apoptotic markers (*casp3b*), retinoic acid metabolism (*cyp26c1*), and tissue hypoxia markers (*hifaa, hifab*). Taken together, this study shows that the *foxe1* mutant zebrafish recapitulates phenotypes associated with FOXE1 mutations in human patients and a clear *foxe1*-ethanol interaction.

## Introduction

Many diseases are rooted in genetic defects, but the expression and severity of symptoms differs between individuals. For example, mutations in genes involved in pathways controlling skeletal development are associated with craniofacial malformations (e.g., cleft palate; (Dixon et al. 2011; Lidral et al. 1998; Park et al. 2007)). At the same time, environmental factors such as drug use, smoking, drinking and other lifestyle habits during pregnancy increase the risk of congenital malformations (van Rooij et al. 2001; DeRoo et al. 2008; Jentink et al. 2010). These findings indicate that genetic factors alone do not fully explain the large variability in skeletal phenotypes in individuals with the same underlying gene mutation. Apparently, it is the interplay between genes and environmental factors that shapes craniofacial development. This interaction is underexplored at present.

During craniofacial development, the transcription factor FOXE1 (Forkhead Box protein e1, also known as thyroid transcription factor 2, TTF2) is involved in proper palate formation and thyroid morphogenesis. Indeed, compromised transcriptional activity of FOXE1, resulting from mutations in the fork head domain of FOXE1, lead to Bamforth-Lazarus syndrome. This syndrome is a rare disease and is characterized by cleft palate and hypothyroidism (Bamforth et al. 1989; Moreno et al. 2009; Castanet and Polak 2010). Single nucleotide polymorphisms located near FOXE1 also significantly affect the risk (both positively and negatively) of developing a cleft palate (Leslie et al. 2017; Imani et al. 2019). How FOXE1-related craniofacial abnormalities arise is not known, and therefore animal models are employed to investigate mechanistic causality between geno- and phenotypes.

Recently, we successfully created a *foxe1* mutant zebrafish that has a compromised Foxe1 function and altered expression of genes controlled by Foxe1 (Raterman et al. 2023). Indeed, in these fish the symptoms of Bamforth-Lazarus syndrome are at least partly recapitulated; affected mutants show aberrant craniofacial development, most notably instability of the ceratohyal elements and decreased bone mineralization. We also detected Foxe1 in the developing thyroid, suggesting conserved function of Foxe1 in zebrafish, but did not observe differences in the anatomy of the developing thyroid. Given that hypothyroidism is a feature of Bamforth-Lazarus syndrome, we addressed the thyroid status of *foxe1* zebrafish mutants in the present study.

Thyroid hormones (TH) regulate a wide array of physiological processes including bone formation and mineralization (Keer et al. 2019) (Bassett and Williams 2016). The activity of the hypothalamus-pituitary-thyroid (HPT) axis is responsible for maintaining normal circulating levels of thyroid hormones. The prohormone thyroxine (or T4) is produced and secreted by the thyroid gland in response to thyroid-stimulating hormone (TSH) from the pituitary gland, which in turn is under control of hypothalamic factors such as, depending on the species, thyrotropin-releasing hormone (TRH) and corticotropin-releasing hormone (CRH) (Ortiga-Carvalho et al. 2016; De Groef et al. 2006; Bernier, Flik, and Klaren 2009). The conversion of T4 to the bioactive 3,3’,5-triiodo-L-thyronine (T3) is catalyzed by iodothyronine deiodinases, types 1 and 2 (Williams and Bassett 2011; Orozco and Valverde-R 2005; Darras 2021). Thyroid hormones bind with nuclear thyroid hormone receptors (TRs) of which different α- and β-isoforms are known (Darras et al. 2011; Ortiga-Carvalho, Sidhaye, and Wondisford 2014). Through their interaction with these receptors, TH modulate gene expression and thereby influence multiple cellular and molecular pathways (Zhang and Lazar 2000).

Another aim of the current study is to understand if interactions between a genetic factor (*foxe1* expression) and an environmental risk factor (ethanol exposure) lead to an abnormally developing craniofacial skeleton. We selected ethanol as environmental risk factor since the craniofacial skeleton is mainly derived from cranial neural crest cells (Knight and Schilling 2006; Mork and Crump 2015; Rocha et al. 2020). By inducing apoptosis of these cells, ethanol induces craniofacial skeletal malformations (Carvan et al. 2004; Lovely et al. 2017). Indeed, sustained maternal ethanol exposure during fetal development causes fetal alcohol spectrum disorders and fetal alcohol syndrome, which both include typical abnormal craniofacial features (Garland, Reynolds, and Zhou 2020). Increased craniofacial sensitivity to ethanol exposure has been described for multiple zebrafish lines with a mutation in the sonic hedgehog (*shh*) pathway, of which *foxe1* is a pathway member (Swartz et al. 2014; Swartz et al. 2020; McCarthy et al. 2013; Everson, Batchu, and Eberhart 2020). In our study, we compared *foxe1* mutant zebrafish to wild types; we investigated mineralization of adult endo- and exoskeletal elements and analyzed the thyroid status. We show that the *foxe1* zebrafish mutant shows features of Bamforth-Lazarus syndrome in adult life stage and that the model offers promising functions as a platform to investigate gene-environment interactions.

## Results

### *foxe1* mutants are phenotypes are suggestive of hypothyroidism

To assess the thyroid status in *foxe1* mutants, we determined the expression of key genes of the HPT-axis. In the pituitary glands of adult mutants, we observed an upregulation of *tshβ*, which is generally accepted to be the primary and most sensitive indicator of (subclinical) hypothyroidism in fish (Figure 1) (Biondi, Cappola, and Cooper 2019). We found no statistically significant differences in *crh* and *trh* expression in the hypothalamus in the mutants compared to wild types.

**Figure 1:**
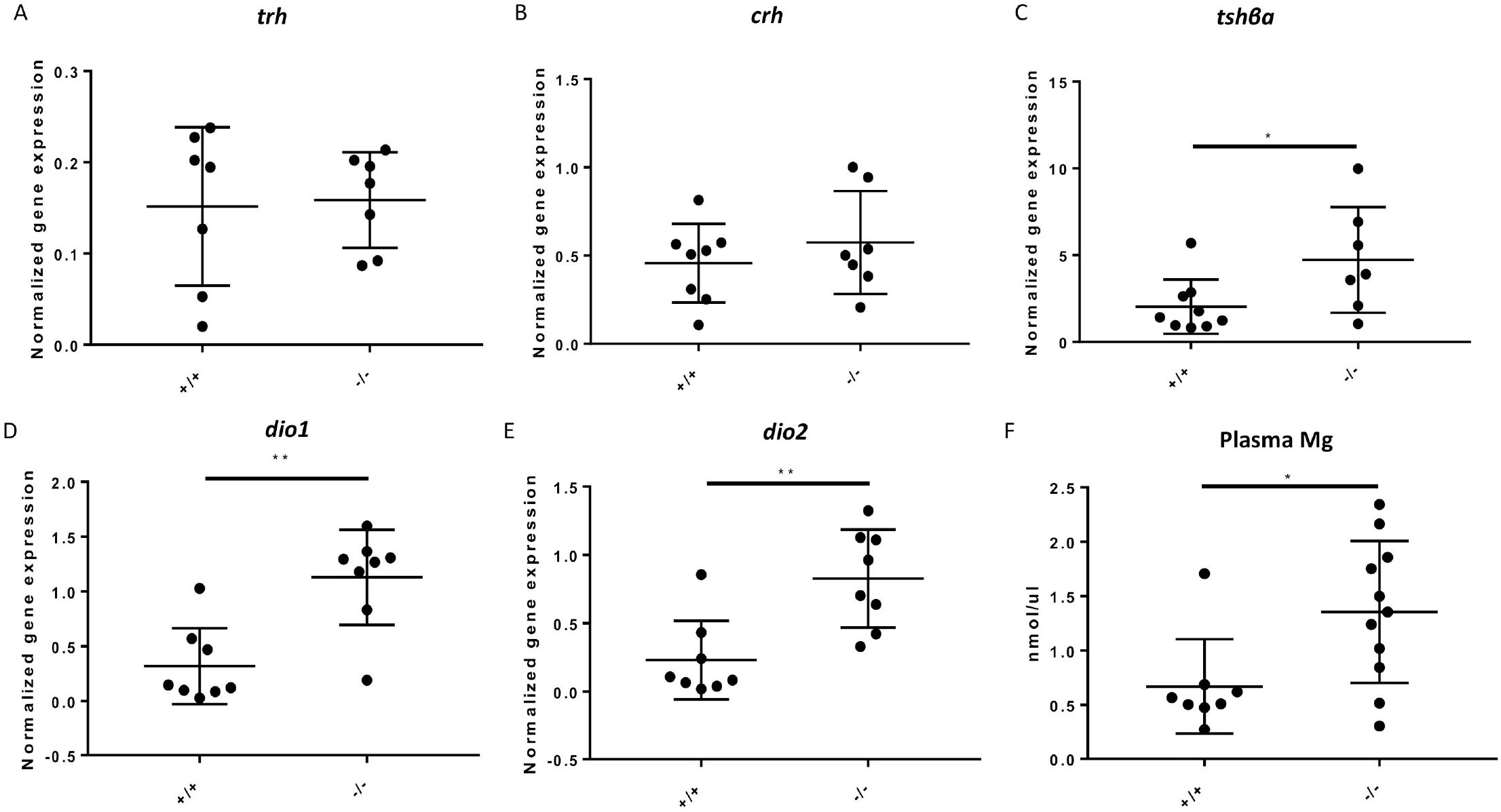
*Foxe1* mutants are hypothyroid. A-B) *trh* and *crh* relative gene expression levels in the hypothalamus of mutants are equal to wild type. C) *tshβ* levels in the pituitary glands of *foxe1*mutants are increased compared to wild types. D-E) Relative expression of deiodinases *dio1* and *dio2* is increased in the livers of mutants compared to wild types (A-E n= 8-10). F) Hypermagnesemia in plasma of *foxe1* mutants (n=19).

Next, we investigated the effects of *foxe1* knockout on peripheral components beyond the HPT-axis. We observed increased relative expressions of *dio1* and *dio2* in the liver of mutants compared to wild types (Figure 1 d, e). This finding further substantiates the notion that *foxe1* mutants are hypothyroid; the same effects were reported for two tilapia species and in striped parrotfish upon methimazole-induced hypothyroidism (Van der Geyten et al. 2001; Johnson and Lema 2011). Moreover, in *foxe1* mutant zebrafish, magnesium plasma concentrations were significantly higher than in wild types (Figure 1f). Magnesium metabolism is consistently affected by hyper- and hypothyroidism in humans; plasma magnesium levels being lowered in hyperthyroidism and elevated in hypothyroidism (Jones et al. 1966).

### *foxe1* mutants have increased osteoclastic activity in scales

Zebrafish scales are part of the dermal skeleton and are easily accessible elements to assess bone metabolism as they harbour all relevant cells involved in bone metabolism (Bergen et al. 2022). A smaller scale size and an altered shape has been reported in several hypothyroid zebrafish mutant models (Aman et al. 2021). Accordingly, in our study we found in scales of *foxe1* mutants a significantly lower molar ratio of calcium:phosphorus (Figure 2a). The mechanical properties of calcium phosphates depend on their calcium:phosphorus ratios (Raynaud et al. 2001) and indeed, lowered ratios in scales of zebrafish have been associated with an osteoporotic phenotype (de Vrieze et al. 2014). Moreover, the scales of *foxe1* mutants were smaller in size and less circular in shape than those of wild types (Figure 2b, c, d). Furthermore, scales of mutants showed increased expression of the osteoclast-specific genes *tracp* and *ctsk*, which are involved in bone resorption and collagen degradation, respectively. The expression of *rankl* was also increased in mutants. RANKL is a ligand produced by osteoblasts which binds to RANK receptors on osteoclast precursors and mature osteoclasts to stimulate fusion and osteoclast activity. Additionally, o*steocalcin* expression was upregulated in *foxe1* mutants. (Figure 2e, f). Tartrate-resistant acid phosphatase (TRAcP) staining confirmed the increased osteoclastic activity in scales of mutants along the radii that we assessed using qPCR (Figure 2i, j). Altogether, these observations indicate an increased activity of osteoclasts in *foxe1* mutant zebrafish.

**Figure 2:**
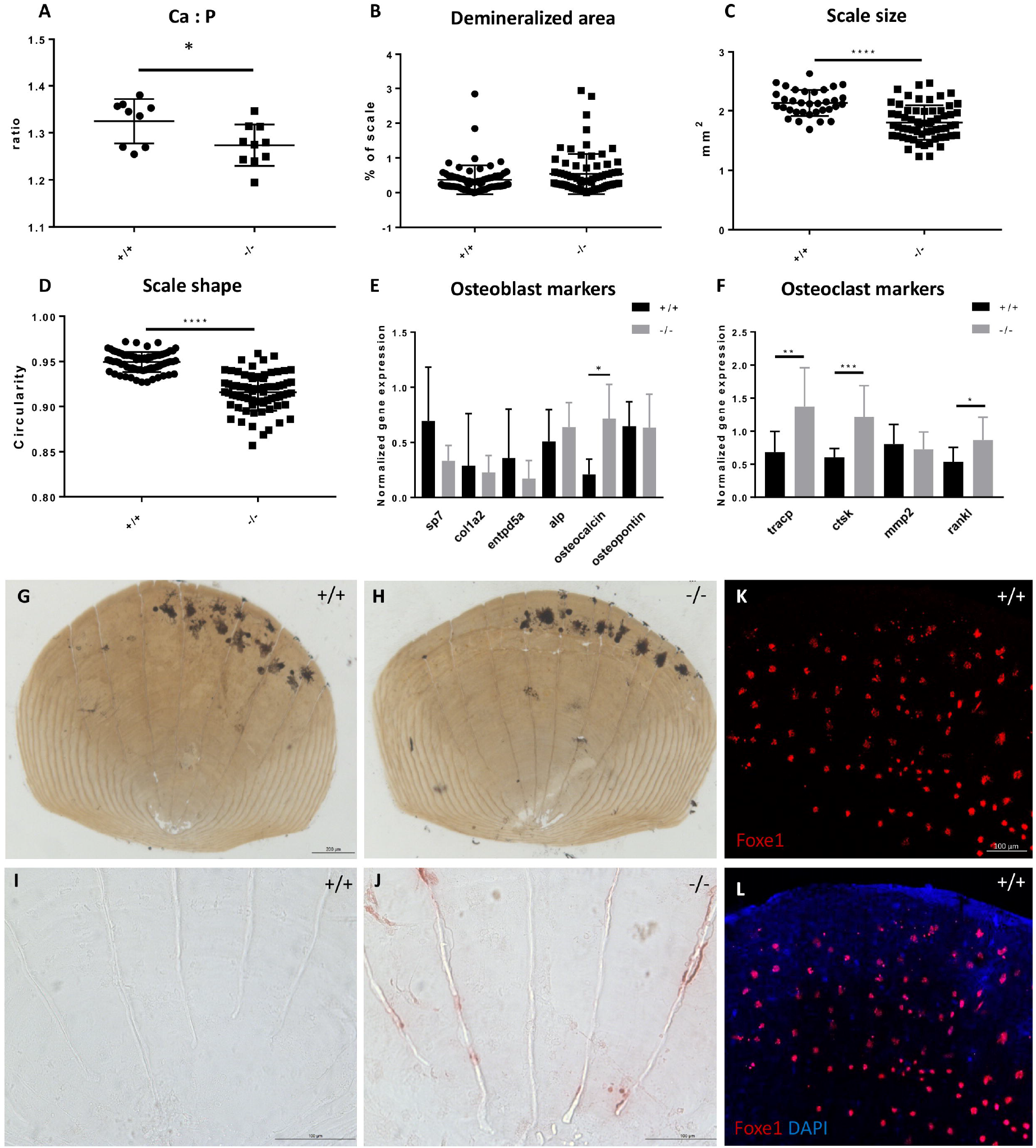
Scales of *foxe1* mutants show increased osteoclast activity and Foxe1 expression. A) The Ca:P ratio of scales is lower in mutants than in *foxe1* wild types. B) The demineralized areas of the scales were not significantly larger in mutants than in wild types. C) The scales of mutants were reduced in size compared to the wild types. D) The shape of the scales of mutants was less round compared to wild types. E) Gene expression of osteoblast markers on wild type and mutant scales. F) Gene expression of osteoclast markers on wild type and mutant scales (n=12). G-H) Wild type and mutant scale stained with Von Kossa to visualize mineralization, unmineralized areas did not differ between mutant and wild type scales, but the shape and size of the scale was affected. I-J) TRAcP staining shows increased osteoclast activity in mutant scales. K-L) *Foxe1*-positive cells at the posterior side of scales in wild type zebrafish, Foxe1 signal and merge.

FOXE1was previously described to be expressed in skin and is involved in the growth and development of skin appendages in different vertebrate lineages (hair, feathers, scales) (Yaklichkin et al. 2011; Han et al. 2018; Eichberger et al. 2004). We indeed found *foxe1*-positive cells in the epidermis at the posterior part of the scale in wild type zebrafish (Figure 2k, l). We asked whether the effects observed in scales of mutants were caused by *foxe1* mutation through altered epidermal-scale interactions or by a change in thyroid status of the mutant. We therefore conducted a TRAcP staining on scales of *dio2*^*-/-*^ zebrafish that are also hypothyroid due to an impaired conversion of the pro-hormone T4 to biologically active T3 (Houbrechts et al. 2016). In *dio2*^*-/-*^ mutants, 81% of mutant scales showed TRAcP-positive cells, while this was 28% in the corresponding controls (Supplemental Figure 1). TRAcP-positive cells were observed in 69% of *foxe1* mutant scales and in 25% of wild type control scales (Figure 2i, j). Thus, in both of the hypothyroid *foxe1* and *dio2* mutants more TRAcP-positive osteoclasts were found compared to wild types.

As larval *foxe1* mutants exhibited reduced mineralization, we also investigated the bone mineralization of adult mutants and wild types. No gross morphological differences were observed after alizarin red staining, nor microCT scanning (Figure 3a-j, Supplemental Figure 2). Also, there was no significant differences in the mineral contents of the lower jaw, the calvaria, the opercle, and the vertebrae between *foxe1* mutants and wild types (Figure 3k). These findings suggest that although *foxe1* mutant larvae are delayed in mineralization, this is no longer seen in adult stages and apparently, they catch up the delay during ontogeny.

**Figure 3:**
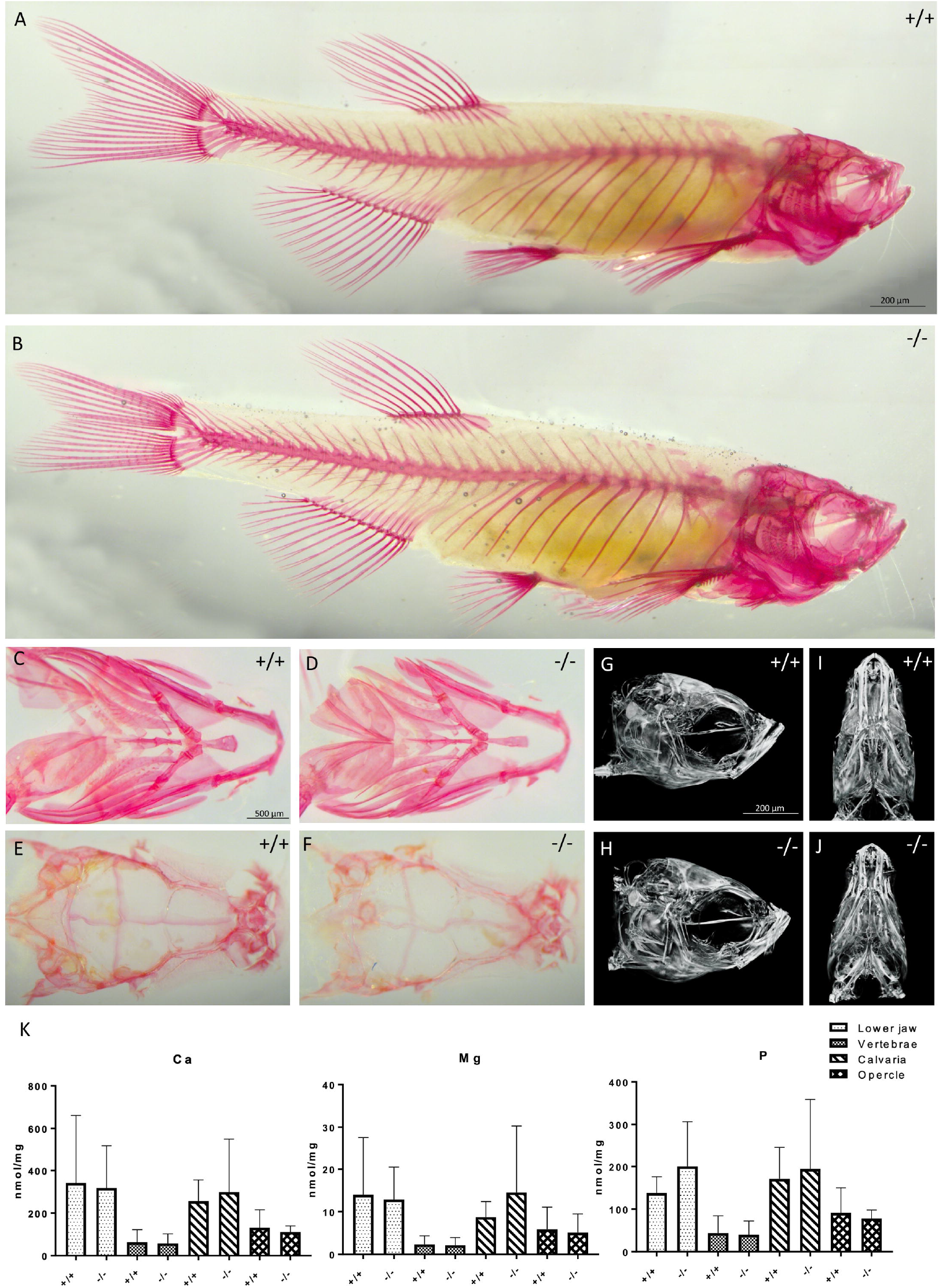
Adult craniofacial skeleton in *foxe1* mutants. A-B) Alizarin red staining of wild type and *foxe1* mutant adult (3 cm SL) zebrafish. C-F) Dissected wild type and mutant lower jaw and calvaria, ventral and dorsal view, respectively (n=13). G-J) 2 cm SL wild type and mutant microCT images (n=3). K) No difference in mineral contents of the lower jaw, vertebrae, calvaria and opercle of wild type and mutant zebrafish (n=10).

### Increased sensitivity to ethanol of craniofacial bone and cartilage in *foxe1* mutants

To investigate whether *foxe1* mutation alters the sensitivity to ethanol, mutant and wild type zebrafish were exposed to varying concentrations of ethanol from 6 to 120 hpf. Subsequently, the development of the skeletal elements was evaluated by bone and cartilage staining (evaluation criteria are described in the methods section). Although malformations were observed in all of the ethanol-exposed groups, the incidence of the malformations was consistently higher in the mutants (Figure 4, all panels).

**Figure 4:**
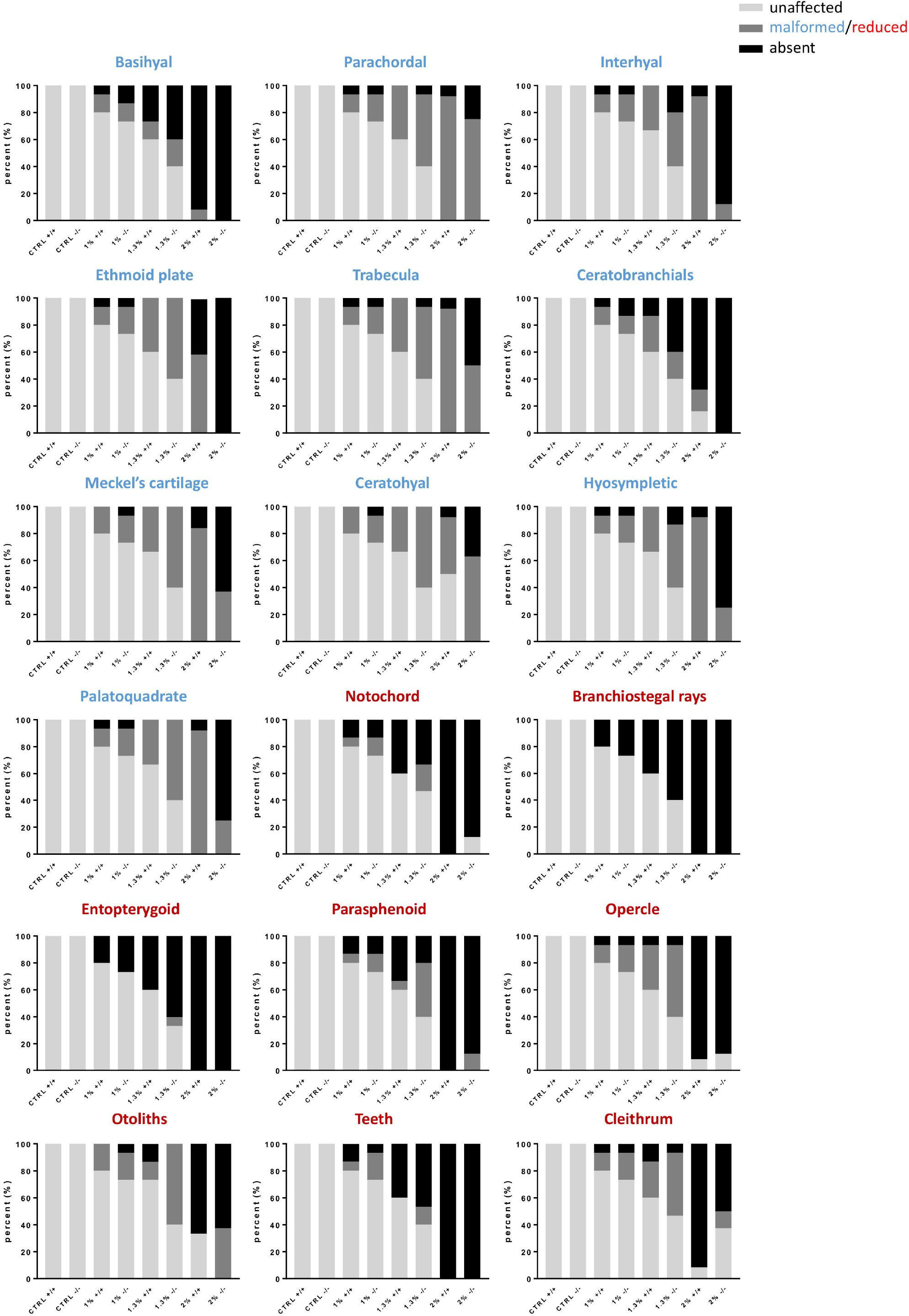
*Foxe1* mutants are more severely affected by ethanol exposure than wild type fish. Wild type and mutant control groups showed normal craniofacial bone and cartilage development. In all other groups, malformations occurred in a subset of the samples. The severity of craniofacial malformations was consistently higher in mutant groups than wild type groups. n =15 for ctrl and 1-1.3% ethanol groups and n=8 for 2% ethanol groups due to poor survival.

We next exposed mutants and wild types to 0% and 1% ethanol and studied craniofacial bone and cartilage defects in more detail. In the control group, no major differences in bone and cartilage formation between wild types and *foxe1* mutants were observed (Figure 5). In the 1% ethanol wild type group, 83% had a straight body form. However, the mineralization of the tip of the notochord, the parasphenoid, the opercles, the cleithrum, the entopterygoid, and the teeth were decreased compared to untreated controls (Figure 5a). We classified this phenotype as mild. Similarly, in the *foxe1* mutant group exposed to 1% ethanol, 72% of the larvae showed a straight body form, but mineralization of the tip of the notochord, the parasphenoid, the opercles, the cleithrum, the entopterygoid and the teeth were decreased compared to untreated mutants (Figure 5a). The effects of 1% ethanol on *foxe1* mutant phenotype were also classified as mild.

**Figure 5:**
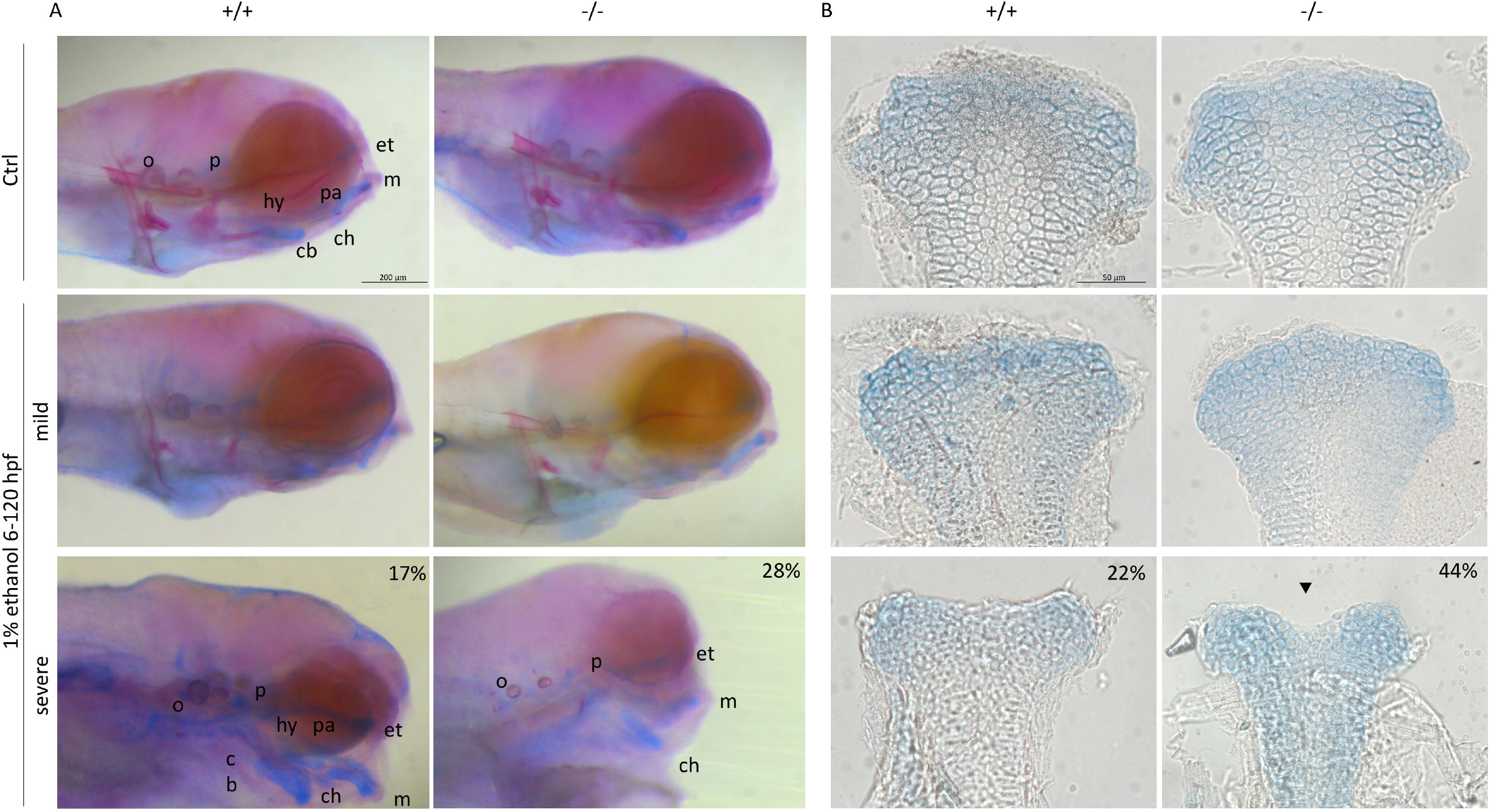
*Foxe1* mutants are severely affected by 1% ethanol exposure. A) Representative images of the effects of 1% ethanol exposure on bone and cartilage development in wild type and mutant larvae at 120 hpf. At 1% ethanol concentration most of the fish were mildly phenotypic, mostly showing defects in bone development. In 17% (6/35) of wild types and 28% (14/49) of mutants more severe phenotypes were observed. In case of a more severe phenotype, the exposed wild types retained more cartilage elements than the exposed mutants. Cb: ceratobranchial, ch: ceratohyal, et: ethmoid plate hy: hyosympletic, m: Meckels, o: otoliths, p: parachordal, pa: palatoquadrate. B) Unexposed control individuals had normal ethmoid plates, while ethanol-exposed larvae had malformed ethmoid plates with dents in the anterior edge or lack of cells in the midline. The incidence was higher in mutants 45/102 (44%) than in wild types 21/92 (22%).

In both mutant and wild type groups, a proportion of ethanol-exposed larvae were severely affected. These larvae had a curved body, a small head, and small eyes. Fewer individuals were severely affected in the wild type group than in the mutant group (17% and 28%, respectively (p=0.22)). In the exposed mutant group, staining revealed fewer bone and cartilage structures than in the exposed wild types: only the otoliths, the parachordal bone, the ethmoid plate, Meckel’s cartilage, and the ceratohyal cartilage were detected (Figure 5a). The number of malformations of cartilage structures was also higher in exposed *foxe1* mutants, and in some cases no neurocranial structures could be distinguished (Figure 5a).

The ethmoid plate is the zebrafish homologue of the human hard palate, and its morphology and growth are affected by ethanol exposure (Duncan et al. 2017; Carvan et al. 2004). We therefore investigated the morphology of this structure in ethanol-exposed larvae and untreated larvae (wild type and mutants). No effects on the morphology of the ethmoid plate were observed in untreated *foxe1* mutants and untreated wild types (Figure 5b). However, following exposure to ethanol, the width and morphology of the ethmoid plates were significantly affected in twice as many mutants than wild types (44% and 22% respectively (p= 0.001)). (Figure 5b). In particular, we observed indentations in the anterior ethmoid plate after ethanol exposure (Figure 5b, arrowhead in lower right panel).

### *foxe1*-ethanol interactions differentially change gene expression

Our next analyses in the investigation of gene-ethanol interactions concerned the expression patterns of marker genes for bone and cartilage development, ethanol clearance, retinoic acid biosynthesis, cell proliferation, and apoptosis. Interactions between the *foxe1* mutant genotype and ethanol exposure significantly affected the expression of the hypoxia-inducible factor zebrafish orthologs *hifaa* and *hifab* (Figure 6a, b) (Supplemental data 1). Moreover, interactions between the mutant genotype and ethanol also significantly affected the expression of the neural crest cell marker *sox9a* as well as *casp3a* and *casp3b*, which play a central role in apoptosis (Figure 6c, d, e) (Supplemental data 1). Upregulation of *casp3b* in exposed mutants indicates increased apoptosis.

**Figure 6:**
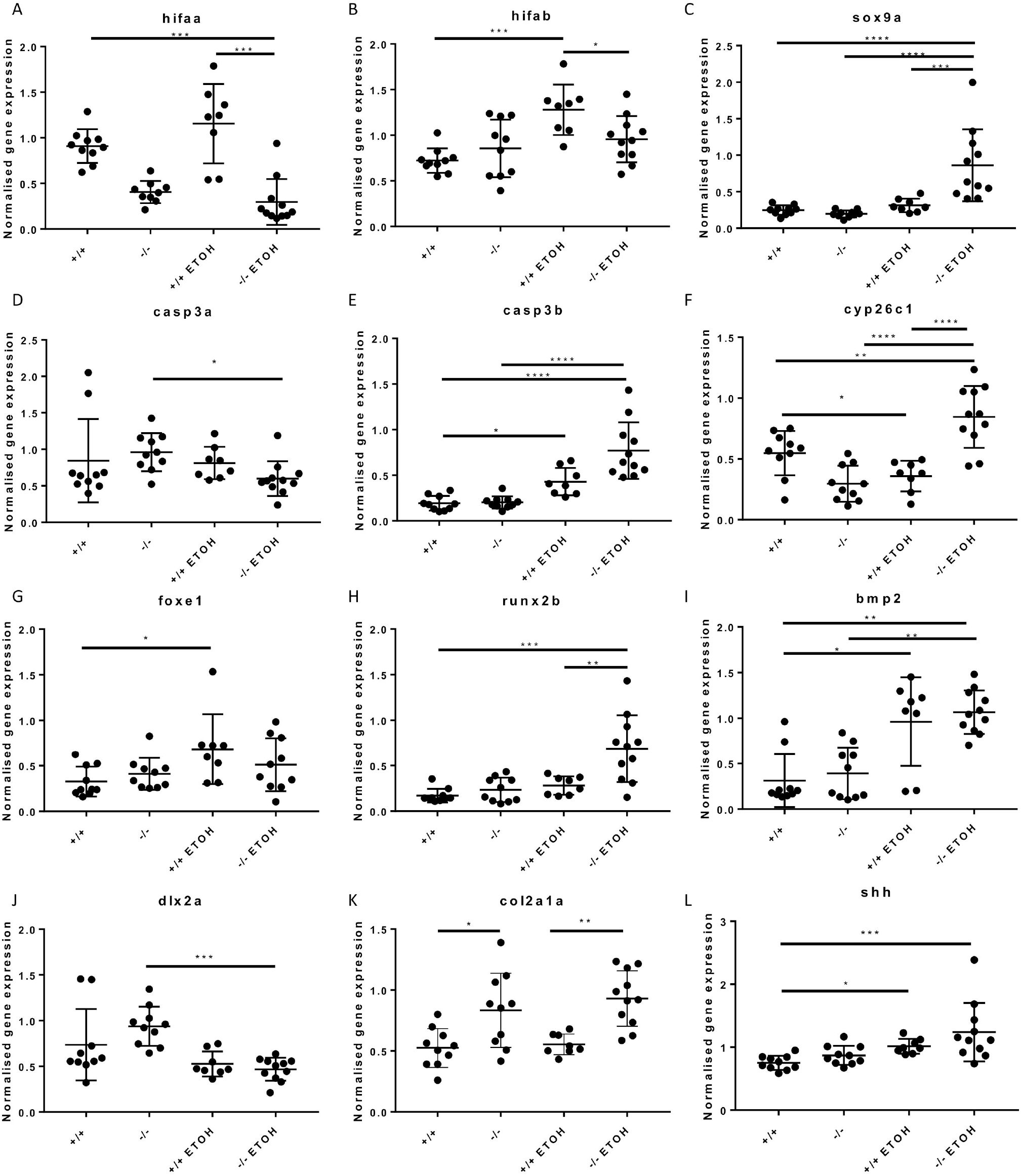
*Foxe1*-ethanol interactions cause differential gene expression of genes involved in cell death, hypoxia, and ethanol clearance. Relative gene expression of *hifaa* (A), *hifab* (B), *sox9a* (C), *casp3a* (D), *casp3b* (E), *cyp26c1* (F), *foxe1* (G), *runx2b* (H), *bmp2* (I), *dlx2a* (J), *col2a1a* (K) and *shh* (L) in *foxe1* mutants and wild types exposed to 0% and 1% ethanol.

Ethanol clearance mechanisms act antagonistically to retinoic acid biosynthesis (Garland, Reynolds, and Zhou 2020; Lovely 2020). Indeed, we found *aldh1a2* and *aldh2*.*2* expression to be upregulated following ethanol exposure, but the *foxe1* genotype did not significantly affect this (Supplemental data 1). We found that gene-ethanol interactions significantly upregulated the expression of *cyp26c1*, which is involved in retinoic acid metabolism (Figure 6f) (Supplemental data 1).

Zhang and colleagues showed previously that transient ethanol exposures in zebrafish result in changed expression of SHH-dependent genes in a zebrafish model of FAS (Zhang et al. 2014). Ethanol exposure of wild type zebrafish indeed upregulated *foxe1* expression (Figure 6g). Interestingly, genes involved in cartilage and bone formation, *runx2b, bmp2, dlx2a, col2a1a*, and *shh* were not significantly affected by *foxe1*-ethanol interactions, but rather by effects of genotype or ethanol separately (Figure 6 h-l) (Supplemental data 1).

## Discussion

In this study we found that *foxe1* zebrafish mutants display characteristics that are distinctive of FOXE1 loss-of-function associated hypothyroidism and skeletal phenotypes in human Bamforth-Lazarus patients. Foxe1 was previously reported to have a key role in the induction and repression of thyroid differentiation and in thyroid hormone synthesis in rat thyroid follicle cells (Lopez-Marquez et al. 2019). The first mouse knockout model of FOXE1-associated hypothyroidism was of only limited value, because knockout was lethal in early development (De Felice et al. 1998). Using a CRE-lox system, Lim and colleagues created knockout Foxe1 mice in adulthood (Lim et al. 2022). The mice became hypothyroid, and this model allowed for studying the role of Foxe1 after thyroid development. In this model, the thyroid follicle morphology changed shortly after knockout, and plasma TSH was upregulated. Both mouse models by De Felice et al. and Lim et al. allowed investigation of the role of Foxe1 in specific life stages exclusively.

In our earlier study, we found FOXE1 in the thyroid follicles of developing zebrafish, which further bolstered the relation between FOXE1 and thyroid metabolism (Raterman et al. 2023). In that study we found no differences in the number of thyroid follicles between mutants and wild type larval fish. The number of follicles varies widely and is not a reliable indicator of thyroid status. In the current study we therefore addressed the thyroid status more thoroughly. Although we were unable to determine circulating levels of thyroid hormones in the plasma (as measurement requires volumes higher than what we can obtain from an adult zebrafish), we see ample evidence to assume that *foxe1* zebrafish mutants are hypothyroid. TSH is widely considered the be the most sensitive indicator of hypothyroidism, e.g. in diagnosis of subclinical cases (Fatourechi 2009). We found a significant upregulation of *tshβ* expression in pituitary glands of adult *foxe1* mutants. Moreover, we observed an upregulation of *dio1* and *dio2* in the liver of *foxe1* mutants, which was also reported three other fish investigations of methimazole-induced hypothyroidism (Van der Geyten et al. 2001; Johnson and Lema 2011). Elevated expression of DIO1 in the liver in mammals would be interpreted as hyperthyroidism because DIO1 expression is positively regulated by thyroid hormone (Maia 2011, Ito 2011). However, it was reported earlier that in zebrafish and other fishes, TH has no effect on *dio1* (Arjona et al. 2011; Walpita, Grommen, et al. 2007; Walpita, Van der Geyten, et al. 2007; Darras 2021). Thirdly, an increase in circulating Mg that we found in the present study, has been linked to thyroid hormone dysfunction (Jones et al. 1966; Simsek et al. 1997).

Thyroid hormones have pervasive effects and target virtually every somatic cell in the body, including skeletal cells. Since *foxe1* is a thyroid transcription factor, we used scales of adult fish to evaluate the effects of the *foxe1* mutation on these skeletal elements. Scale formation is initiated in the skin, which itself is a direct target of TH as transporters and (co)receptors of TH are expressed in the skin (Aman et al. 2021). In a zebrafish model of hypothyroidism, *thyroglobulin* was ablated in larvae at 5 dpf. Later in ontogeny, scale development in these fish was severely affected. Compared to controls, the hypothyroid fish developed twice as many scales, which were miniaturized and disorderly arranged. The onset of scale formation in these mutants was delayed until the fish were much larger, leading to a higher number of smaller scales. Loss of *dio2* activity led to the same phenotype (Aman et al. 2021). Starting from these notions, we investigated the scales in our mutant. Consistent with the two models of hypothyroidism, *foxe1* mutants show an altered scale shape and size. Yet, the *foxe1* mutant scale phenotype is less dramatic, suggesting either the activation of compensatory mechanisms or residual function of the hypomorph allele.

In mammals, hypothyroidism leads to delayed endochondral ossification, impaired growth, and reduced mineralization (Devdhar, Ousman, and Burman 2007; Bassett and Williams 2016; Persani 2012). Previously we reported that skeletal mineralization during endochondral ossification is decreased in *foxe1* mutant larvae (Raterman et al. 2023), but at that time we could not relate this to hypothyroidism yet. We now assessed a reduced mineralization and an increased osteoclast activity in scales from *foxe1* and *dio2* zebrafish mutants which we link to hypothyroidism. An increase in osteoclast activity is typically associated with human hyperthyroidism, but accelerated bone remodeling, reduced bone density, and osteoporosis are also described (Reddy et al. 2012). The symptoms of hypothyroidism are usually the opposite of those of hyperthyroidism and thyrotoxicosis, but there is evidence that this is not the case for bone metabolism. For example, fracture risk increases in both hyper- and hypothyroid patients (Vestergaard and Mosekilde 2002). Also, in children with congenital hypothyroidism bone mineral density is decreased compared to controls (Demartini Ade et al. 2007).

The risk of developing congenital malformations such as orofacial clefts is heavily impacted by environmental factors and gene-environment interactions (van Rooij et al. 2001; DeRoo et al. 2008; Jentink et al. 2010). Specific gene polymorphisms that confer sensitivity together with environmental exposure further increase the risk. For instance, risks of developing craniofacial malformations are increased by maternal smoking and drinking in combination with specific variants of *TGFB3* and *MSX1* (Romitti et al. 1999). Similarly, polymorphisms in *MTHFR* increase the risk of developing CL/P, which can be attenuated by taking supplemental folate (Ibarra-Lopez et al. 2013). Assessing the risks of gene specific interactions and generalized effects of environmental factors can contribute to lifestyle recommendations for pregnant women. However, research in that field of gene-environment interactions is still limited. Until now, only one interaction of FOXE1 with an environmental factor was described; two FOXE1 polymorphisms increased susceptibility to metabolic and thyroid dysfunction caused by nitrates in drinking water (Gandarilla-Esparza et al. 2021). Apart from a useful model for hypothyroidism, the *foxe1* mutant zebrafish also appears to be suitable for gene-environment interaction studies to craniofacial malformations.

Mutations and polymorphisms in *shh* pathway were previously described to increase sensitivity to ethanol (Swartz et al. 2014; Swartz et al. 2020; McCarthy et al. 2013; Kietzman et al. 2014; Everson, Batchu, and Eberhart 2020). To determine whether the *foxe1* mutation also influences the sensitivity to ethanol, we exposed developing zebrafish to various ethanol concentrations. Indeed, ethanol exposure affected the mutants more severely than the wild types. In all tested concentrations, ethanol increased the risk of skeletal malformations as well as the severity of the malformation. Both bone and cartilage structures were malformed or even absent more often in the mutants than in the wild types upon ethanol exposure. In the *foxe1* ethanol treated mutants, all elements of the head skeleton were affected, but the neurocranium structures, including the ethmoid plate, were affected more severely. The ethmoid plate is regarded as an analogue of the human palate (Duncan et al. 2017). Specifically, the width of the ethmoid plate was decreased, while the anterior ethmoid was irregularly shaped. Various specific gene-ethanol interactions have been described earlier in the development of the ethmoid plate. When *pdgfra* mutants were exposed to ethanol, no ethmoid plate was formed at all, while hypoplasia of other craniofacial structures was also described (McCarthy et al. 2013). McCarthy and colleagues reported that 20% of the wild type embryos exposed to ethanol developed severe craniofacial defects, similar to the wild types in the present study (McCarthy et al. 2013). *Vangl2* mutants exposed to ethanol form only a small rod instead of a full ethmoid plate, while *mars* mutants develop gaps in the posterior ethmoid plate (Swartz et al. 2014). This variation in malformations in different mutants following ethanol exposure highlights the specificity of gene-ethanol interactions. Moreover, loss-of-function mutations in a number of important craniofacial genes such as *cyp26b1, gata3, smoothened* and *smad5* did not modify the risk for ethanol-induced craniofacial defects in zebrafish (Smith et al. 2014; McCarthy et al. 2013). This further emphasizes that the increased effects of ethanol in sensitive mutants are caused by specific gene-ethanol interactions.

The effects of ethanol on early development are pleiotropic as effects on cell death, cell proliferation, retinoic acid signaling, and oxidative stress have been described (Fish et al. 2021; Fernandes and Lovely 2021; Garland, Reynolds, and Zhou 2020). Previous work showed that the loss of tp53 protects against ethanol-induced craniofacial malformations, illustrating that apoptosis is a pathogenic mechanism of ethanol teratogenesis (Fish et al. 2021). In *foxe1* mutant, *tp53* signaling was not affected by gene-ethanol interaction (Supplemental data 1), but *caspase* expression had increased. Moreover, hypoxia-inducible factor gene expression had also decreased in the exposed mutants, which suggests that the mutant is less capable of responding to oxidative stress upon ethanol exposure. Hypoxia was previously shown to cause clefts of the anterior ethmoid plate in zebrafish and increased cleft palate incidence in mice (Kuchler et al. 2018; Millicovsky and Johnston 1981).

Our data thus point toward an increased sensitivity of the *foxe1* mutant to ethanol. We used previously described concentrations and regimes for ethanol exposure. At 1% (∼160 mM) ethanol, tissue levels in zebrafish reach about 41mM (0.16 %).(McCarthy et al. 2013; Swartz et al. 2014). This is consistent with concentrations that can be reached in humans during binge drinking events (Lange and Voas 2000). However, the exposure duration used in the experiments may have magnified the effects of ethanol on development as zebrafish were exposed throughout embryonic and larval development (Fernandes and Lovely 2021).

This study set out to investigate the effects of a *foxe1* mutation in zebrafish. In humans, FOXE1 mutation causes Bamforth-Lazarus syndrome. We found a phenotype similar to hypothyroidism in adult mutants and the consequences on scale formation and squamation. Moreover, we found evidence of increased sensitivity of *foxe1* mutants to ethanol in larvae leading to more, and severely affected craniofacial malformations. Altogether, this provides insight into the role of *FOXE1* in non-syndromic craniofacial malformations. Subtle mutations or gene variations that do not give rise to pronounced malformations alone may cause isolated craniofacial malformations upon exposure to environmental factors such as ethanol. The versatile zebrafish model can be used further to investigate the role of FOXE1 in thyroid development and to screen for possible gene-environment interactions in the etiology of craniofacial malformations.

## Methods

### Animals

Animal procedures were carried out in accordance with the Dutch Animals Act and European laws. Ethical approval for the experiments was granted by Radboud University’s Institutional Animal Care and Use Committee (AVD10300202115245). Zebrafish *foxe1^rdb2* mutants (Raterman et al. 2023) and wild types were kept at standard care conditions (28°C under a 14 h light/10 h dark cycle) in the Radboud University Zebrafish Facility. Embryos and larvae were raised in petri dishes with E3 medium (5 mM NaCl, 0.17 mM KCl, 0.33 mM CaCl_2_, 0.33 mM MgSO_4_, 0.00001% Methylene Blue) in an incubator at 28.5°C with a 14 h light/10 h dark cycle. For ethanol interaction studies, larvae were exposed to 1% (v/v) ethanol in E3 medium between 6 hpf and 120 hpf.

### Bone staining of adult zebrafish

Bone staining was performed as described by Sakata-Haga and colleagues (Sakata-Haga et al. 2018). *foxe1^rdb2* mutants and wild types with a standard length of 2 cm (approximately 2.5 months old) were selected and euthanized using 2-phenoxyethanol, fixed and cleared using 5% formalin, 5% Triton X-100, 1% potassium hydroxide (KOH) at 42°C. Then the samples were cleared further in 20% ethylene glycol, 5% Triton X-100, 1% KOH at 42°C. Next, the samples were pre-stained using 20% ethylene glycol, 1% KOH before bone staining using 0.05% alizarin red S, 20% ethylene glycol, 1% KOH. Samples were then descaled and cleared in 20% Tween 20, 1% KOH at 42°C. Samples were then transferred to 99% glycerol for imaging on a binocular microscope using Leica Application Suite (LAS 3.3, Leica).

### Inductively Coupled Plasma - Optical Emission Spectroscopy

Tissues and blood were collected, tissues were dried and weighed, and then 100 μL of 65% nitric acid (HNO_3_) was added. Nitric acid containing the dissolved samples was added to 6 mL ultrapure water. Inductively Coupled Plasma - Optical Emission Spectroscopy (ICP-OES) was used to determine the molar contents of calcium, phosphorus, and magnesium in the samples. Samples were measured on an ARCOSMV (Spectro, Kleve, Germany) on axial view, with 1400 W plasma power. Samples were nebulized with a SeaSpray nebulizer combined with a cyclone chamber and an argon flow of 0.7 L/min.

### Micro-CT analyses

*foxe1^rdb2* mutants and wild types with a standard length of 2 cm were euthanized using 2-phenoxyethanol and fixed in 4% PFA. The samples were scanned on a Bruker MicroCT Skyscan 1172 (Bruker). For micro-CT scanning the X-ray source was set to 60 kV and 156 μA. Rotation step 0.43□ at 360□, with a frame average of 4. Pixel size was 6.0 μm. No filter was used. Images were rendered using Skyscan and CTX software.

### Von Kossa staining

Scales were collected from a standardized area on the left flank of the wild type and *foxe1* mutant zebrafish directly after euthanizing. The scales were rinsed in demineralized water before incubation in 5% silver nitrate solution. The staining was fixed using a 5% sodium thiosulfate solution. Scales were imaged on binocular microscope using Leica Application Suite (LAS 3.3, Leica). Scales were assessed for size, demineralized area and circularity using the ZF macro (Tarasco et al. 2020).

### TRAcP staining

Scales were collected from a standardized area on the left flank directly after euthanizing. The scales were preincubated in 0.2 M Tris buffer at 37□C. After rinsing in distilled water the TRAcP staining was performed as described by (van de Wijngaert and Burger 1986) at 37 □C. The scales were imaged on a binocular microscope using Leica Application Suite (LAS 3.3, Leica).

### Immunohistochemistry

Whole mount immunohistochemistry was performed on the scales exactly as described previously (Raterman et al. 2023) using a Foxe1 antibody (Boster Bio (DZ41149), Pleasanton, USA) at 1:500. Z-stack images of the T4 staining were taken on an SPX8 confocal microscope (Leica).

### qPCR

Total larval RNA of 5 dpf *foxe1^rdb2* mutants and wild type zebrafish, and dissected adult tissues were extracted in 400 μl Trizol reagent (Invitrogen, Carlsbad, USA) following the manufacturer’s protocol. Yields were quantified using a NANODROP 1000 spectrophotometer (manufacturer, city, country). cDNA was synthesized using the iScript cDNA Synthesis Kit (Bio-Rad, Cressier, Switzerland). The master mix consisted of 10 μl SYBR green mix (2X) (Bio-Rad, Hercules, USA) and qPCR was performed using a CFX 96 (Bio-Rad) machine. Primers are listed in table 1. Relative gene expression was determined after normalization against the reference genes *elf1a* and *rpl13*. For pituitary tissue, *fshβ* was used as a tissue-specific housekeeping gene (Vandesompele et al. 2002). Larval data were analyzed with a two-way ANOVA on raw or transformed data. Tissue data were analyzed with a Student’s *t*-test (in case of two groups) or two-way ANOVA or Kruskal-Wallis test (in case of more than two groups), where appropriate.

**Table 1.**
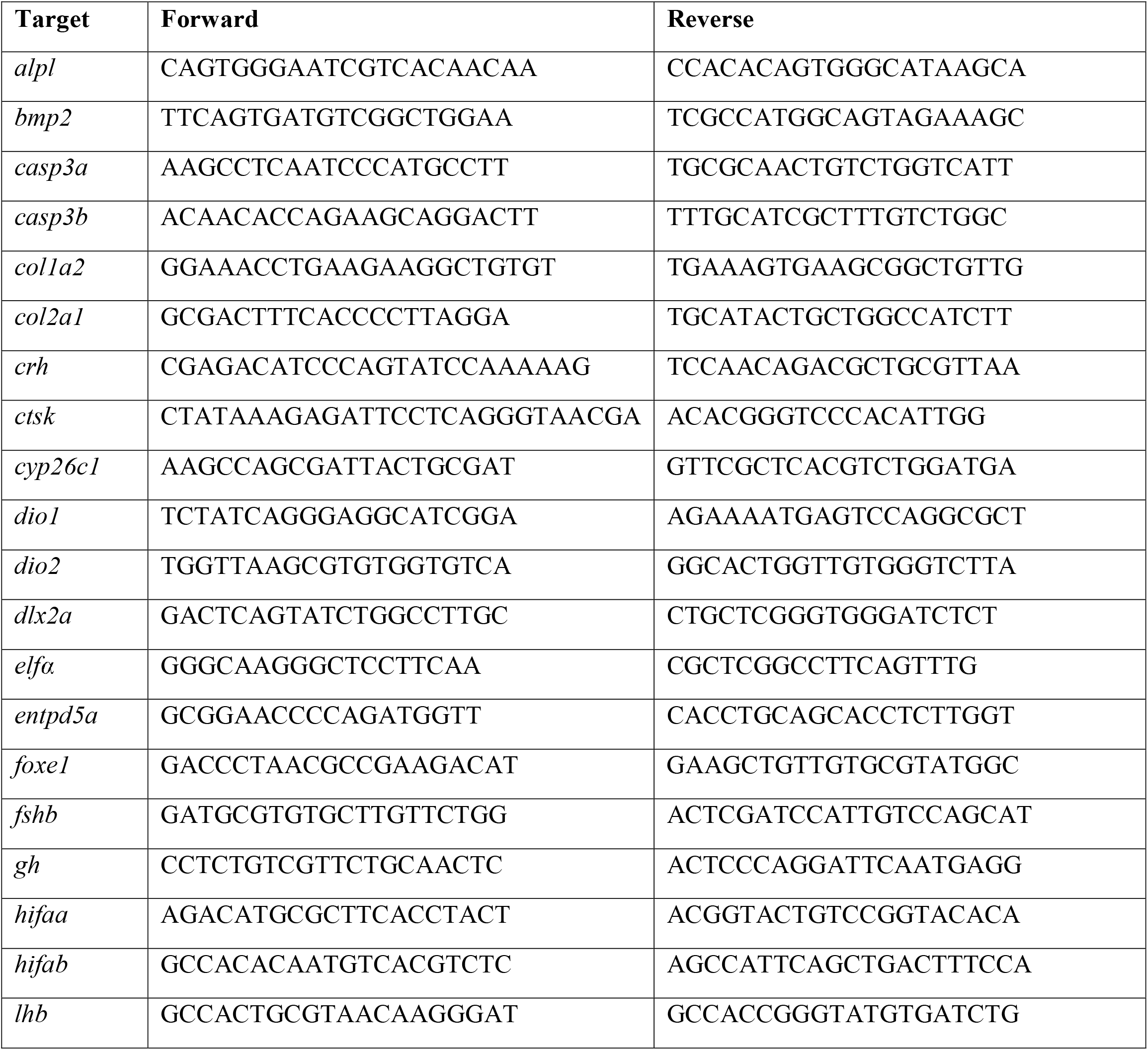
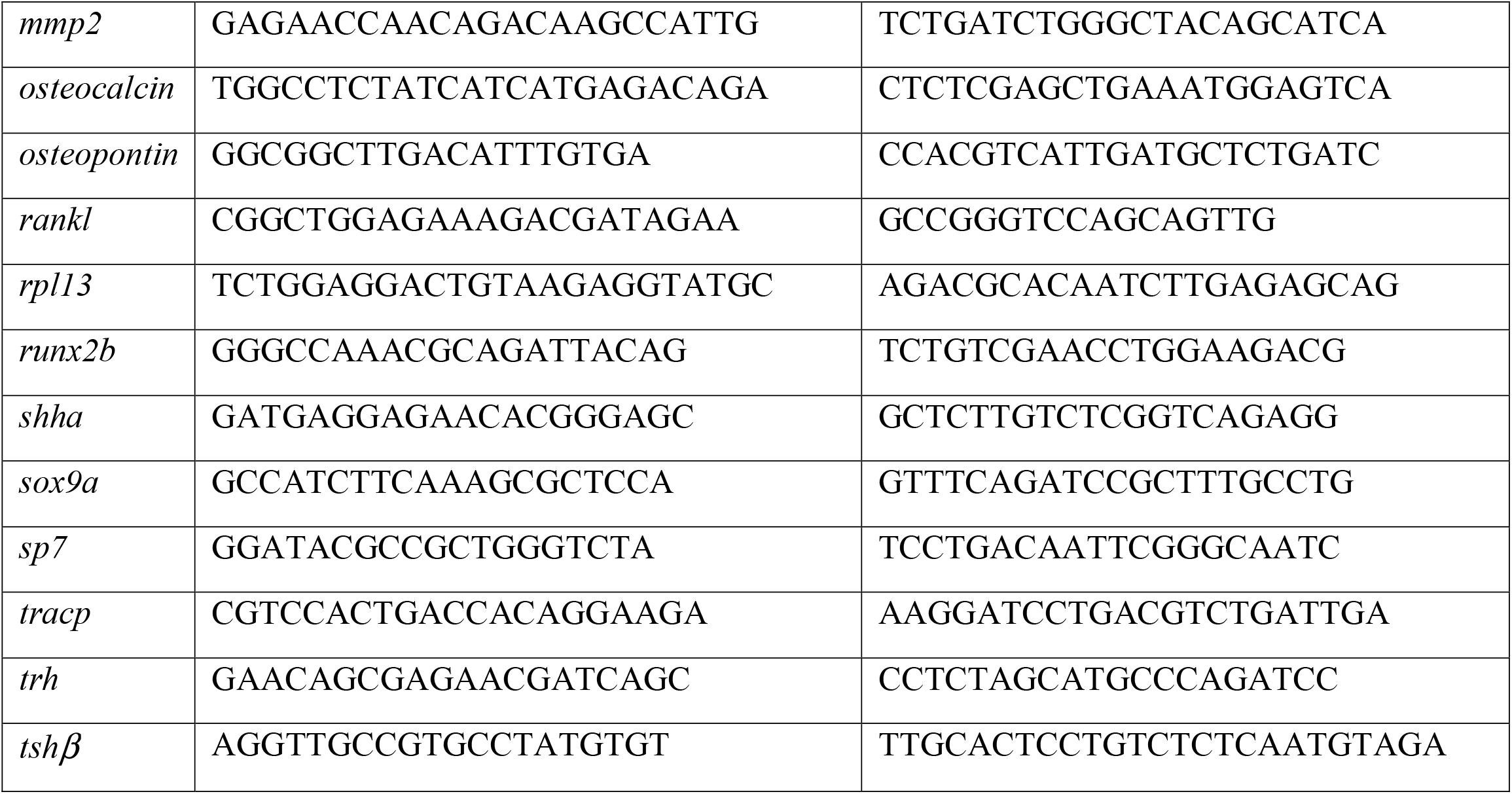
qPCR primer sequences.

### Alizarin red and Alcian blue staining of larvae

Staining was performed as descried by Walker and Kimmel (Walker and Kimmel 2007). In brief, after fixation for 1 hour in 2% PFA a 100 mM Tris/40 mM MgCl_2_ (pH 7.5) wash was performed. Then the cartilage staining solution consisting of 0.04% Alcian Blue (Sigma-Aldrich) in 40 mM MgCl_2_ was applied for 2.5 h. After gradual rehydration, samples were bleached using 3% H_2_O_2_ in 0.5% KOH. After two washes in 25% glycerol/0.1% KOH, bone staining was performed in 0.03% Alizarin Red (pH 7.5, Sigma-Aldrich) for 1 hour. Next, larvae were imaged in 3% methylcellulose on a binocular microscope using Leica Application Suite (LAS 3.3, Leica). After imaging, the ethmoid plates were dissected and mounted on glass slides for further detailed imaging.

For figure 4, larvae were assessed by eye and scored for the formation of craniofacial skeletal elements. Larvae were exposed to 1-2% ethanol in the E3 medium from 6 hpf until 120 hpf and stained for bone and cartilage at 120 hpf. Then, the craniofacial skeleton was screened for the development of each bone or cartilage element. Animals were scored to have normal, malformed, or completely absent cartilage structures (graphs with a blue heading). Animals were scored to have normally mineralized, reduced mineralized, or completely absent bone structures (graphs with a red heading). Data were analyzed with a Chi-square test.

## Acknowledgements

We thank Antoon van der Horst and Jeroen Boerrigter for animal care at the Radboud University Zebrafish Facility. Paul van de Ven and Sebastian Krosse are gratefully acknowledged for the elemental analyses.

## Data availability

All relevant data can be found within the article and its supplementary information. Further inquiries can be directed to the corresponding author.

## Figure legends

Supplemental figure 1: **TRAcP staining in *dio2* mutants**. In *dio2* mutants 81% of mutant scales showed TRAcP staining and in the corresponding controls 28% of scales showed staining (n=20-38). Dio2 mutants were generated and characterized by (Houbrechts et al. 2016).

Supplemental figure 2: **Alizarin red staining on *foxe1* mutants at 2 and 3 cm standard length**

## References

Aman, A. J., M. Kim, L. M. Saunders, and D. M. Parichy. 2021. ‘Thyroid hormone regulates abrupt skin morphogenesis during zebrafish postembryonic development’, Developmental Biology, 477: 205–18.

Arjona, F. J., L. Vargas-Chacoff, M. P. Martin Del Rio, G. Flik, J. M. Mancera, and P. H. Klaren. 2011. ‘Effects of cortisol and thyroid hormone on peripheral outer ring deiodination and osmoregulatory parameters in the Senegalese sole (Solea senegalensis)’, J Endocrinol, 208: 323–30.

Bamforth, J. S., I. A. Hughes, J. H. Lazarus, C. M. Weaver, and P. S. Harper. 1989. ‘Congenital hypothyroidism, spiky hair, and cleft palate’, Journal of Medical Genetics, 26: 49–51.

Bassett, J. H., and G. R. Williams. 2016. ‘Role of Thyroid Hormones in Skeletal Development and Bone Maintenance’, Endocr Rev, 37: 135–87.

Bergen, D. J. M., Q. Tong, A. Shukla, E. Newham, J. Zethof, M. Lundberg, R. Ryan, S. E. Youlten, M. Frysz, P. I. Croucher, G. Flik, R. J. Richardson, J. P. Kemp, C. L. Hammond, and J. R. Metz. 2022. ‘Regenerating zebrafish scales express a subset of evolutionary conserved genes involved in human skeletal disease’, BMC Biol, 20: 21.

Bernier, N. J., G. Flik, and P. H. M. Klaren. 2009. ‘Regulation and Contribution of the Corticotropic, Melanotropic and Thyrotropic Axes to the Stress Response in Fishes’, Fish Neuroendocrinology, 28: 235–311.

Biondi, B., A. R. Cappola, and D. S. Cooper. 2019. ‘Subclinical Hypothyroidism: A Review’, JAMA, 322: 153–60.

Carvan, M. J., 3rd, E. Loucks, D. N. Weber, and F. E. Williams. 2004. ‘Ethanol effects on the developing zebrafish: neurobehavior and skeletal morphogenesis’, Neurotoxicol Teratol, 26: 757–68.

Castanet, M., and M. Polak. 2010. ‘Spectrum of Human Foxe1/TTF2 Mutations’, Hormone Research in Paediatrics, 73: 423–29.

Darras, V. M. 2021. ‘Deiodinases: How Nonmammalian Research Helped Shape Our Present View’, Endocrinology, 162.

Darras, V. M., S. L. Van Herck, M. Heijlen, and B. De Groef. 2011. ‘Thyroid hormone receptors in two model species for vertebrate embryonic development: chicken and zebrafish’, J Thyroid Res, 2011: 402320.

De Felice, M., C. Ovitt, E. Biffali, A. Rodriguez-Mallon, C. Arra, K. Anastassiadis, P. E. Macchia, M. G. Mattei, A. Mariano, H. Scholer, V. Macchia, and R. Di Lauro. 1998. ‘A mouse model for hereditary thyroid dysgenesis and cleft palate’, Nature Genetics, 19: 395–8.

De Groef, B., S. Van der Geyten, V. M. Darras, and E. R. Kuhn. 2006. ‘Role of corticotropin-releasing hormone as a thyrotropin-releasing factor in non-mammalian vertebrates’, General and Comparative Endocrinology, 146: 62–68.

de Vrieze, E., M. A. van Kessel, H. M. Peters, F. A. Spanings, G. Flik, and J. R. Metz. 2014. ‘Prednisolone induces osteoporosis-like phenotype in regenerating zebrafish scales’, Osteoporos Int, 25: 567–78.

Demartini Ade, A., C. A. Kulak, V. C. Borba, M. N. Cat, R. S. Dondoni, R. Sandrini, S. Nesi-Franca, and Ld Lacerda Filho. 2007. ‘[Bone mineral density of children and adolescents with congenital hypothyroidism]’, Arq Bras Endocrinol Metabol, 51: 1084–92.

DeRoo, L. A., A. J. Wilcox, C. A. Drevon, and R. T. Lie. 2008. ‘First-trimester maternal alcohol consumption and the risk of infant oral clefts in Norway: a population-based case-control study’, American Journal of Epidemiology, 168: 638–46.

Devdhar, M., Y. H. Ousman, and K. D. Burman. 2007. ‘Hypothyroidism’, Endocrinol Metab Clin North Am, 36: 595–615, v.

Dixon, M. J., M. L. Marazita, T. H. Beaty, and J. C. Murray. 2011. ‘Cleft lip and palate: understanding genetic and environmental influences’, Nat Rev Genet, 12: 167–78.

Duncan, K. M., K. Mukherjee, R. A. Cornell, and E. C. Liao. 2017. ‘Zebrafish models of orofacial clefts’, Dev Dyn, 246: 897–914.

Eichberger, T., G. Regl, M. S. Ikram, G. W. Neill, M. P. Philpott, F. Aberger, and A. M. Frischauf. 2004. ‘FOXE1, a new transcriptional target of GLI2 is expressed in human epidermis and basal cell carcinoma’, J Invest Dermatol, 122: 1180–7.

Everson, J. L., R. Batchu, and J. K. Eberhart. 2020. ‘Multifactorial Genetic and Environmental Hedgehog Pathway Disruption Sensitizes Embryos to Alcohol-Induced Craniofacial Defects’, Alcohol Clin Exp Res, 44: 1988–96.

Fatourechi, V. 2009. ‘Subclinical hypothyroidism: an update for primary care physicians’, Mayo Clin Proc, 84: 65–71.

Fernandes, Y., and C. B. Lovely. 2021. ‘Zebrafish models of fetal alcohol spectrum disorders’, Genesis, 59: e23460.

Fish, E. W., S. K. Tucker, R. L. Peterson, J. K. Eberhart, and S. E. Parnell. 2021. ‘Loss of tumor protein 53 protects against alcohol-induced facial malformations in mice and zebrafish’, Alcohol Clin Exp Res, 45: 1965–79.

Gandarilla-Esparza, D. D., E. Y. Calleros-Rincon, H. M. Macias, M. F. Gonzalez-Delgado, G. G. Vargas, J. D. Sustaita, A. Gonzalez-Zamora, E. Rios-Sanchez, and R. Perez-Morales. 2021. ‘FOXE1 polymorphisms and chronic exposure to nitrates in drinking water cause metabolic dysfunction, thyroid abnormalities, and genotoxic damage in women’, Genet Mol Biol, 44: e20210020.

Garland, M. A., K. Reynolds, and C. J. Zhou. 2020. ‘Environmental mechanisms of orofacial clefts’, Birth Defects Res, 112: 1660–98.

Han, W., X. Li, L. Wang, H. Wang, K. Yang, Z. Wang, R. Wang, R. Su, Z. Liu, Y. Zhao, Y. Zhang, and J. Li. 2018. ‘Expression of fox-related genes in the skin follicles of Inner Mongolia cashmere goat’, Asian-Australas J Anim Sci, 31: 316–26.

Houbrechts, A. M., J. Delarue, I. J. Gabriels, J. Sourbron, and V. M. Darras. 2016. ‘Permanent Deiodinase Type 2 Deficiency Strongly Perturbs Zebrafish Development, Growth, and Fertility’, Endocrinology, 157: 3668–81.

Ibarra-Lopez, J. J., P. Duarte, V. Antonio-Vejar, E. S. Calderon-Aranda, G. Huerta-Beristain, E. Flores-Alfaro, and M. E. Moreno-Godinez. 2013. ‘Maternal C677T MTHFR polymorphism and environmental factors are associated with cleft lip and palate in a Mexican population’, J Investig Med, 61: 1030–5.

Imani, M. M., M. Safaei, P. Lopez-Jornet, and M. Sadeghi. 2019. ‘A systematic review and meta-analysis on protective role of forkhead box E1 (FOXE1) polymorphisms in susceptibility to non-syndromic cleft lip/palate’, Int Orthod, 17: 437–45.

Jentink, J., M. A. Loane, H. Dolk, I. Barisic, E. Garne, J. K. Morris, L. T. de Jong-van den Berg, and Eurocat Antiepileptic Study Working Group. 2010. ‘Valproic acid monotherapy in pregnancy and major congenital malformations’, N Engl J Med, 362: 2185–93.

Johnson, K. M., and S. C. Lema. 2011. ‘Tissue-specific thyroid hormone regulation of gene transcripts encoding iodothyronine deiodinases and thyroid hormone receptors in striped parrotfish (Scarus iseri)’, Gen Comp Endocrinol, 172: 505–17.

Jones, J. E., P. C. Desper, S. R. Shane, and E. B. Flink. 1966. ‘Magnesium metabolism in hyperthyroidism and hypothyroidism’, Journal of Clinical Investigation, 45: 891–900.

Keer, S., K. Cohen, C. May, Y. N. Hu, S. McMenamin, and L. P. Hernandez. 2019. ‘Anatomical Assessment of the Adult Skeleton of Zebrafish Reared Under Different Thyroid Hormone Profiles’, Anatomical Record-Advances in Integrative Anatomy and Evolutionary Biology, 302: 1754–69.

Kietzman, H. W., J. L. Everson, K. K. Sulik, and R. J. Lipinski. 2014. ‘The teratogenic effects of prenatal ethanol exposure are exacerbated by Sonic Hedgehog or GLI2 haploinsufficiency in the mouse’, Plos One, 9: e89448.

Knight, R. D., and T. F. Schilling. 2006. ‘Cranial neural crest and development of the head skeleton’, Adv Exp Med Biol, 589: 120–33.

Kuchler, E. C., L. A. D. Silva, P. Nelson-Filho, T. M. Saboia, A. M. Rentschler, J. M. Granjeiro, D. Oliveira, P. N. Tannure, R. A. D. Silva, L. S. Antunes, M. Tsang, and A. R. Vieira. 2018. ‘Assessing the association between hypoxia during craniofacial development and oral clefts’, J Appl Oral Sci, 26: e20170234.

Lange, J. E., and R. B. Voas. 2000. ‘Defining binge drinking quantities through resulting BACs’, Annu Proc Assoc Adv Automot Med, 44: 389–404.

Leslie, E. J., J. C. Carlson, J. R. Shaffer, A. Butali, C. J. Buxo, E. E. Castilla, K. Christensen, F. W. Deleyiannis, L. Leigh Field, J. T. Hecht, L. Moreno, I. M. Orioli, C. Padilla, A. R. Vieira, G. L. Wehby, E. Feingold, S. M. Weinberg, J. C. Murray, T. H. Beaty, and M. L. Marazita. 2017. ‘Genome-wide meta-analyses of nonsyndromic orofacial clefts identify novel associations between FOXE1 and all orofacial clefts, and TP63 and cleft lip with or without cleft palate’, Hum Genet, 136: 275–86.

Lidral, A. C., P. A. Romitti, A. M. Basart, T. Doetschman, N. J. Leysens, S. Daack-Hirsch, E. V. Semina, L. R. Johnson, J. Machida, A. Burds, T. J. Parnell, J. L. Rubenstein, and J. C. Murray. 1998. ‘Association of MSX1 and TGFB3 with nonsyndromic clefting in humans’, American Journal of Human Genetics, 63: 557–68.

Lim, G., A. Widiapradja, S. P. Levick, K. J. McKelvey, X. H. Liao, S. Refetoff, M. Bullock, and R. J. Clifton-Bligh. 2022. ‘Foxe1 Deletion in the Adult Mouse Is Associated With Increased Thyroidal Mast Cells and Hypothyroidism’, Endocrinology, 163.

Lopez-Marquez, A., C. Fernandez-Mendez, P. Recacha, and P. Santisteban. 2019. ‘Regulation of Foxe1 by Thyrotropin and Transforming Growth Factor Beta Depends on the Interplay Between Thyroid-Specific, CREB and SMAD Transcription Factors’, Thyroid, 29: 714–25.

Lovely, C. B. 2020. ‘Animal models of gene-alcohol interactions’, Birth Defects Res, 112: 367–79.

Lovely, C., M. Rampersad, Y. Fernandes, and J. Eberhart. 2017. ‘Gene-environment interactions in development and disease’, Wiley Interdiscip Rev Dev Biol, 6.

McCarthy, N., L. Wetherill, C. B. Lovely, M. E. Swartz, T. M. Foroud, and J. K. Eberhart. 2013. ‘Pdgfra protects against ethanol-induced craniofacial defects in a zebrafish model of FASD’, Development, 140: 3254–65.

Millicovsky, G., and M. C. Johnston. 1981. ‘Hyperoxia and hypoxia in pregnancy: simple experimental manipulation alters the incidence of cleft lip and palate in CL/Fr mice’, Proc Natl Acad Sci U S A, 78: 5722–3.

Moreno, L. M., M. A. Mansilla, S. A. Bullard, M. E. Cooper, T. D. Busch, J. Machida, M. K. Johnson, D. Brauer, K. Krahn, S. Daack-Hirsch, J. L’Heureux, C. Valencia-Ramirez, D. Rivera, A. M. Lopez, M. A. Moreno, A. Hing, E. J. Lammer, M. Jones, K. Christensen, R. T. Lie, A. Jugessur, A. J. Wilcox, P. Chines, E. Pugh, K. Doheny, M. Arcos-Burgos, M. L. Marazita, J. C. Murray, and A. C. Lidral. 2009. ‘FOXE1 association with both isolated cleft lip with or without cleft palate, and isolated cleft palate’, Human Molecular Genetics, 18: 4879–96.

Mork, L., and G. Crump. 2015. ‘Zebrafish Craniofacial Development: A Window into Early Patterning’, Curr Top Dev Biol, 115: 235–69.

Orozco, A., and C. Valverde-R. 2005. ‘Thyroid hormone deiodination in fish’, Thyroid, 15: 799–813.

Ortiga-Carvalho, T. M., M. I. Chiamolera, C. C. Pazos-Moura, and F. E. Wondisford. 2016. ‘Hypothalamus-Pituitary-Thyroid Axis’, Compr Physiol, 6: 1387–428.

Ortiga-Carvalho, T. M., A. R. Sidhaye, and F. E. Wondisford. 2014. ‘Thyroid hormone receptors and resistance to thyroid hormone disorders’, Nat Rev Endocrinol, 10: 582–91.

Park, J. W., I. McIntosh, J. B. Hetmanski, E. W. Jabs, C. A. Vander Kolk, Y. H. Wu-Chou, P. K. Chen, S. S. Chong, V. Yeow, S. H. Jee, B. Y. Park, M. D. Fallin, R. Ingersoll, A. F. Scott, and T. H. Beaty. 2007. ‘Association between IRF6 and nonsyndromic cleft lip with or without cleft palate in four populations’, Genet Med, 9: 219–27.

Persani, L. 2012. ‘Clinical review: Central hypothyroidism: pathogenic, diagnostic, and therapeutic challenges’, J Clin Endocrinol Metab, 97: 3068–78.

Raterman, S. T., J. W. Von Den Hoff, S. Dijkstra, C. De Vriend, T. Te Morsche, S. Broekman, J. Zethof, E. De Vrieze, Fadtg Wagener, and J. R. Metz. 2023. ‘Disruption of the foxe1 gene in zebrafish reveals conserved functions in development of the craniofacial skeleton and the thyroid’, Front Cell Dev Biol, 11: 1143844.

Raynaud, S., E. Champion, D. Bernache-Assollant, and J. P. Laval. 2001. ‘Determination of calcium/phosphorus atomic ratio of calcium phosphate apatites using X-ray diffractometry’, Journal of the American Ceramic Society, 84: 359–66.

Reddy, P. A., C. V. Harinarayan, A. Sachan, V. Suresh, and G. Rajagopal. 2012. ‘Bone disease in thyrotoxicosis’, Indian J Med Res, 135: 277–86.

Rocha, M., N. Singh, K. Ahsan, A. Beiriger, and V. E. Prince. 2020. ‘Neural crest development: insights from the zebrafish’, Dev Dyn, 249: 88–111.

Romitti, P. A., A. C. Lidral, R. G. Munger, S. Daack-Hirsch, T. L. Burns, and J. C. Murray. 1999. ‘Candidate genes for nonsyndromic cleft lip and palate and maternal cigarette smoking and alcohol consumption: evaluation of genotype-environment interactions from a population-based case-control study of orofacial clefts’, Teratology, 59: 39–50.

Sakata-Haga, H., M. Uchishiba, H. Shimada, T. Tsukada, M. Mitani, T. Arikawa, H. Shoji, and T. Hatta. 2018. ‘A rapid and nondestructive protocol for whole-mount bone staining of small fish and Xenopus’, Sci Rep, 8: 7453.

Simsek, G., G. Andican, Y. Karakoc, G. Yigit, H. Hatemi, and G. Candan. 1997. ‘Calcium, magnesium, and zinc status in experimental hypothyroidism’, Biol Trace Elem Res, 60: 205–13.

Smith, S. M., A. Garic, M. E. Berres, and G. R. Flentke. 2014. ‘Genomic factors that shape craniofacial outcome and neural crest vulnerability in FASD’, Front Genet, 5: 224.

Swartz, M. E., C. B. Lovely, N. McCarthy, T. Kuka, and J. K. Eberhart. 2020. ‘Novel Ethanol-Sensitive Mutants Identified in an F3 Forward Genetic Screen’, Alcohol Clin Exp Res, 44: 56–65.

Swartz, M. E., M. B. Wells, M. Griffin, N. McCarthy, C. B. Lovely, P. McGurk, J. Rozacky, and J. K. Eberhart. 2014. ‘A screen of zebrafish mutants identifies ethanol-sensitive genetic loci’, Alcohol Clin Exp Res, 38: 694–703.

Tarasco, M., F. P. Cordelieres, M. L. Cancela, and V. Laize. 2020. ‘ZFBONE: An ImageJ toolset for semi-automatic analysis of zebrafish bone structures’, Bone, 138: 115480.

van de Wijngaert, F. P., and E. H. Burger. 1986. ‘Demonstration of tartrate-resistant acid phosphatase in un-decalcified, glycolmethacrylate-embedded mouse bone: a possible marker for (pre)osteoclast identification’, J Histochem Cytochem, 34: 1317–23.

Van der Geyten, S., A. Toguyeni, J. F. Baroiller, B. Fauconneau, A. Fostier, J. P. Sanders, T. J. Visser, E. R. Kuhn, and V. M. Darras. 2001. ‘Hypothyroidism induces type I iodothyronine deiodinase expression in tilapia liver’, Gen Comp Endocrinol, 124: 333–42.

van Rooij, I. A., M. J. Wegerif, H. M. Roelofs, W. H. Peters, A. M. Kuijpers-Jagtman, G. A. Zielhuis, H. M. Merkus, and R. P. Steegers-Theunissen. 2001. ‘Smoking, genetic polymorphisms in biotransformation enzymes, and nonsyndromic oral clefting: a gene-environment interaction’, Epidemiology, 12: 502–7.

Vandesompele, J., K. De Preter, F. Pattyn, B. Poppe, N. Van Roy, A. De Paepe, and F. Speleman. 2002. ‘Accurate normalization of real-time quantitative RT-PCR data by geometric averaging of multiple internal control genes’, Genome Biology, 3.

Vestergaard, P., and L. Mosekilde. 2002. ‘Fractures in patients with hyperthyroidism and hypothyroidism: a nationwide follow-up study in 16,249 patients’, Thyroid, 12: 411–9.

Walker, M. B., and C. B. Kimmel. 2007. ‘A two-color acid-free cartilage and bone stain for zebrafish larvae’, Biotechnic & Histochemistry, 82: 23–28.

Walpita, C. N., S. V. Grommen, V. M. Darras, and S. Van der Geyten. 2007. ‘The influence of stress on thyroid hormone production and peripheral deiodination in the Nile tilapia (Oreochromis niloticus)’, Gen Comp Endocrinol, 150: 18–25.

Walpita, C. N., S. Van der Geyten, E. Rurangwa, and V. M. Darras. 2007. ‘The effect of 3,5,3’-triiodothyronine supplementation on zebrafish (Danio rerio) embryonic development and expression of iodothyronine deiodinases and thyroid hormone receptors’, Gen Comp Endocrinol, 152: 206–14.

Williams, G. R., and J. H. Bassett. 2011. ‘Deiodinases: the balance of thyroid hormone: local control of thyroid hormone action: role of type 2 deiodinase’, J Endocrinol, 209: 261–72.

Yaklichkin, S. Y., D. K. Darnell, M. V. Pier, P. B. Antin, and S. Hannenhalli. 2011. ‘Accelerated evolution of 3’avian FOXE1 genes, and thyroid and feather specific expression of chicken FoxE1’, Bmc Evolutionary Biology, 11: 302.

Zhang, C., J. M. Frazier, H. Chen, Y. Liu, J. A. Lee, and G. J. Cole. 2014. ‘Molecular and morphological changes in zebrafish following transient ethanol exposure during defined developmental stages’, Neurotoxicol Teratol, 44: 70–80.

Zhang, J., and M. A. Lazar. 2000. ‘The mechanism of action of thyroid hormones’, Annu Rev Physiol, 62: 439–66.

