## Supplementary figures and images for "*foxe1* mutant zebrafish show indications of a hypothyroid phenotype and increased sensitivity to ethanol for craniofacial malformations"

### Supplemental figure 1

Supplemental figure 1

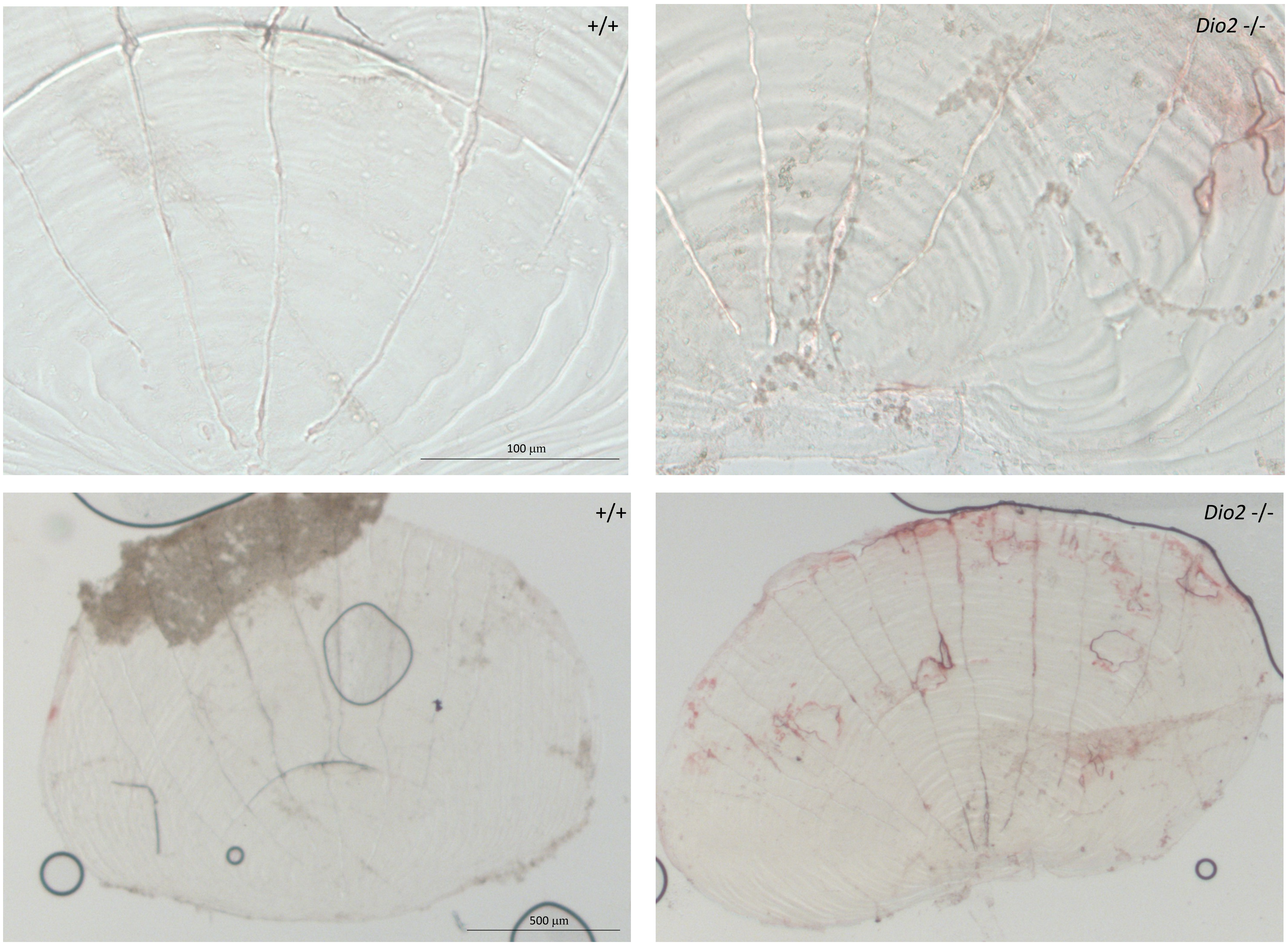

### Supplemental figure 2

Supplemental figure 2

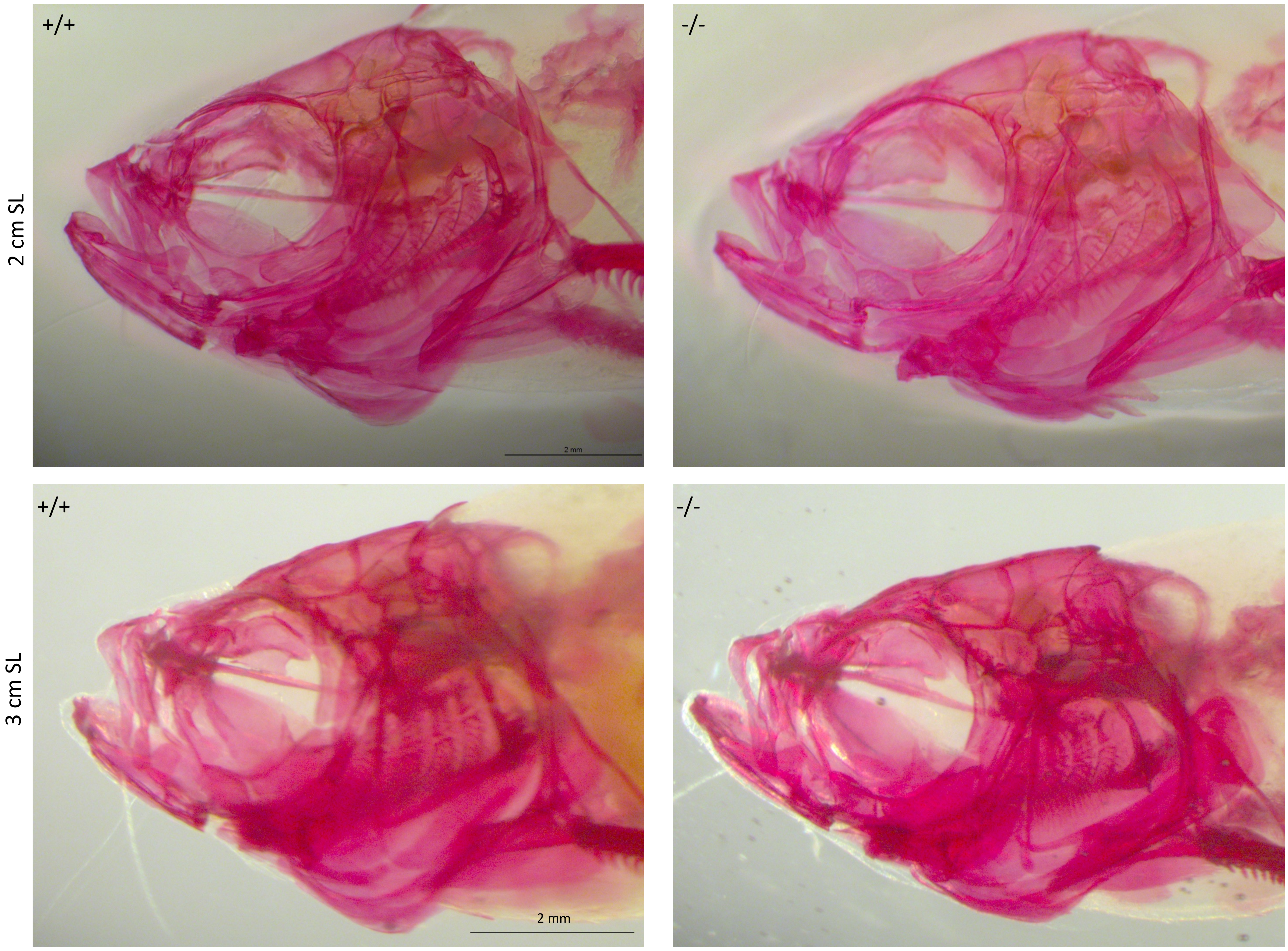
